# Cortical areas for planning sequences before and during movement

**DOI:** 10.1101/2023.11.05.565682

**Authors:** Giacomo Ariani, Mahdiyar Shahbazi, Jörn Diedrichsen

## Abstract

Production of rapid movement sequences relies on preparation before (pre-planning) and during (online planning) movement. Here, we asked how different cortical sensorimotor areas contribute to these processes. Human participants performed three single-finger and three multi-finger sequences in a delayed movement paradigm. During preparation, 7T functional MRI revealed that primary motor (M1) and somatosensory (S1) areas showed pre-activation of the first movement, even though the overall activation level did not change from baseline. During production, the activity in M1 and S1 could be explained by temporal summation of activity patterns corresponding to constituent fingers. In contrast, dorsal premotor (PMd) and anterior superior parietal lobule (aSPL) showed substantial activation during preparation of multi-finger as compared to single-finger sequences. The same regions were also more activated during production of multi-finger sequences, suggesting that the same areas are involved in both pre- and online planning. Nonetheless, we observed small but robust differences between the two contrasts, suggesting preferential involvements of these areas in pre- and online planning. Multivariate analysis revealed sequence-specific representations in both PMd and aSPL, which remained stable across both preparation and production phases. This suggests that these areas maintain a sequence-specific representation before and during sequence production, likely guiding the execution-related areas.

**Significance Statement:** Understanding how the brain orchestrates complex behavior remains a core challenge in human neuroscience. Here, we combine high-resolution neuroimaging and a carefully crafted design to study the neural control of rapid sequential finger movements, like typing or playing the piano. Advancing prior research, we show that the brain areas involved in planning these movements maintain those representations throughout the execution of the sequence. This representational stability across planning and execution suggests an intricate connection between these processes. Our results shed light on the nuanced contributions of different cortical areas to different aspects of coordinating skilled movement. This work is well placed to inform future research in animal models and the development of targeted interventions against movement disorders.

## Introduction

From buttoning a shirt to texting with a smartphone, many everyday actions depend on the brain’s ability to coordinate rapid sequences of finger movements. Behavioral studies have demonstrated that, when tasked to produce a sequence of multiple finger presses, participants pre-plan the first two or three elements before sequence production starts (Ariani & Diedrichsen, 2019; Ariani et al., 2021). Once the sequence starts, planning the upcoming movements continues throughout sequence production, a process called online planning. Most behavioral improvements during motor sequence learning can be explained by participants becoming faster at pre- and online planning (Ariani & Diedrichsen, 2019). Which, and how, different cortical motor areas contribute to these different aspects of motor sequence planning, however, is poorly understood.

Previous neuroimaging studies of motor sequences have used multivariate analysis of fMRI data to reveal a hierarchy of sequence representations across cortical motor areas (Berlot et al., 2020; Yokoi et al., 2018; Yokoi & Diedrichsen, 2019). The dorsal premotor cortex (PMd) and the superior parietal lobule (SPL) exhibit sequence-specific representations, i.e., activity patterns that encode the specific sequence of actions, not just the individual movements themselves. Sequence-specific representations in association cortex have also been shown using electrophysiology in non-human primates (Russo et al., 2020; Shima et al., 2006; Tanji & Shima, 1994). In contrast, activity patterns in the primary motor (M1) and somatosensory cortex (S1) could be explained by a summation of the patterns related to the individual finger presses (Berlot et al., 2021; Yokoi et al., 2018). Due to difficulties related to fMRI temporal resolution, however, this work did not distinguish between activity arising from sequence planning or execution. Although Gallivan et al. (2016) showed that sequences of two upper-limb actions (e.g., reaching to grasp and place vs. hold a cup) could be distinguished from preparatory fMRI activity patterns, but the nature of these representations remains unknown.

Here, we used high-field (7T) fMRI while human participants planned and executed both multi- and single-finger sequences on a keyboard device (matched number of keypresses across sequences). A delayed-movement paradigm with no-go trials (Ariani et al., 2018) allowed us to isolate brain activity related to planning and execution and address the following three questions about the role of cortical areas in sequence planning.

First, we investigated brain responses in primary sensorimotor cortex (M1 and S1) during the preparation of a sequence. For single-finger actions, we have previously shown that movement planning pre-activates the relevant finger-specific activity pattern in both M1 and S1 (Ariani et al., 2022). But what is the preparatory state in M1 for multi-finger movements?

Second, recent behavioral studies suggest that online planning of movement sequences shares important behavioral features with sequence pre-planning—both exhibit a similar planning horizon and both contribute to performance improvements with sequence learning (Ariani & Diedrichsen, 2019; Ariani et al., 2021). We therefore tested to what degree pre- and online planning engage the same cortical areas by contrasting multi- and single-finger movements during preparation and production phases.

Finally, employed multi-variate analysis to study two of the identified brain regions, PMd and SPL, in more depth. Previous results (Berlot et al., 2021; Yokoi & Diedrichsen, 2019) have revealed sequence-specific representations in both. Our paradigm now allowed us to ask whether these representations are present only during preparation or whether they persist during movement production. Furthermore, we investigated to what degree the sequence-specific representations underlying pre- and online planning are the same, or whether they dynamically change from preparation to production.

## Materials and Methods

### Participants

Twenty-three right-handed neurologically healthy participants volunteered to take part in the experiment (13 F, 10 M; age 20–31 years, mean 23.43 years, SD 4.08 years). Criteria for inclusion were right-handedness and no prior history of psychiatric or neurological disorders. Handedness was assessed with the Edinburgh Handedness Inventory (mean 82.83, SD 9.75). All experimental procedures were approved by the Research Ethics Committee at Western University. Participants provided written informed consent to procedures and data usage and received monetary compensation for their participation. One participant withdrew before study completion and was excluded from data analysis (final N = 22). A part of the data used in the current paper was previously published (Ariani et al., 2022).

### Apparatus

Sequences of right-hand finger presses were performed on a custom-made MRI-compatible keyboard device (Fig. 1A). Participants used their fingers to press the keys. The keys of the device did not move, but force transducers underneath each key measured isometric force production at an update rate of 2 ms (Honeywell FS series; dynamic range 0-25 N). A keypress/release was detected when the force crossed a threshold of 1 N. The forces measured from the keyboard were low pass filtered to reduce noise induced by the MRI environment, amplified, and sent to the PC for online task control and data recording.

**Figure 1.**
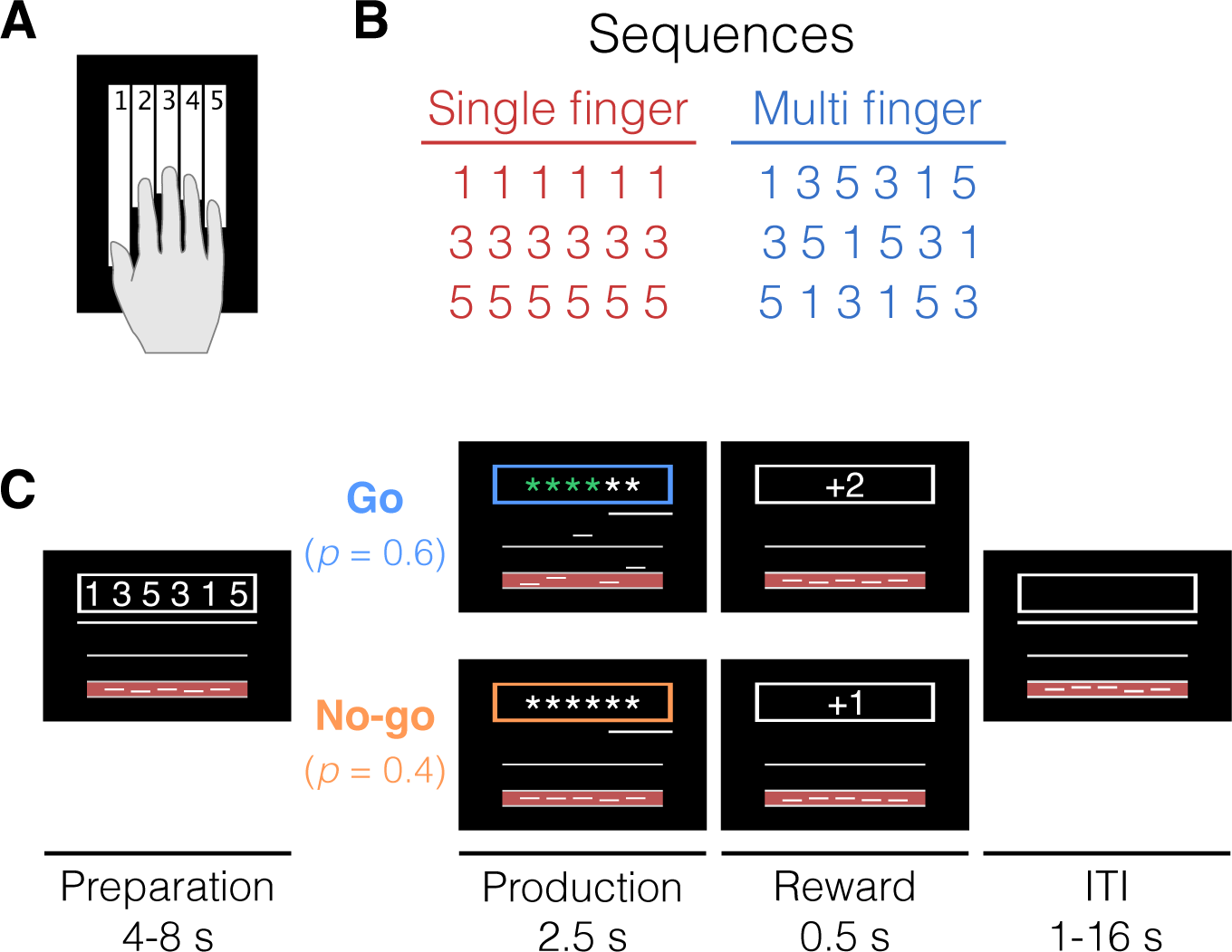
Sequence conditions and task timing. **(A)** Response keyboard with mapping between numbers and fingers. Numbers were not visible on the keys. **(B)** Digits presented on the screen for single-finger (red) and multi-finger sequences (blue). **(C)** Temporal structure of a trial. During the preparation phase, a sequence of 6 numbers was displayed within a box at the top of the screen. With a probability of 0.6, the box frame then changed to a blue color, instructing participants (N=22) to produce the memorized sequence as fast as possible (go trials). Each correct press caused an asterisk to turn green. With 0.4 probability, the box frame turned orange, signaling participants to withhold the response (no-go trials). ITI = inter-trial interval.

### Task

We used a task in which participants produced sequences of keypresses with their right-hand fingers in response to numerical cues (Fig. 1B) presented on a computer screen that was visible to the participants lying in the scanner through an angled mirror. On each trial, a string of 6 numbers (instructing cue) instructed which sequence to plan (Fig. 1C, white outline).

The length of the preparation phase was randomly sampled to be 4 s (56% of trials), 6 s (30%), or 8 s (14%). To control for involuntary overt movements during the preparation phase, we required participants to maintain a steady force of around 0.25 N on all the keys during the delay, which was closely monitored online. As a visual aid, we displayed a red area (between 0 and 0.5 N) and asked participants to remain in the middle of that range with all the fingers (touching either boundary of the red area would count as unwanted movement, thus incurring an error).

At the onset of the production phase, participants received a color cue (go/no-go cue) indicating whether to perform the planned finger presses (blue outline = go, p = 0.6) or not (orange outline = no-go, p = 0.4). The role of no-go trials was to dissociate the hemodynamic response to the successive preparation and production events, which would otherwise always overlap in fast fMRI designs due to the sluggishness of the fMRI response. To encourage planning during the delay period, at the go cue, the digits were masked with asterisks at go-cue onset, and participants had to perform the presses from memory. Participants had 2.5 s to complete the sequence of 6 presses, and a vanishing white bar under the asterisks indicated how much time was left. Participants received online feedback on the correctness of each press with asterisks turning either green, for a correct press or red, for incorrect presses. As long as the participants remained within task constraints (i.e., 6 keypresses in less than 2.5 s), an exact movement speed was not enforced. In no-go trials, participants were instructed to remain as still as possible, maintaining the finger pre-activation until the end of the production phase (i.e., releasing any of the keys would incur an error).

During the reward phase (0.5 s), points were awarded based on performance and according to the following scheme: -1 point in case of no-go error or go cue anticipation (timing errors); 0 points for pressing any wrong key (press error); 1 point in case of a correct no-go trial; and 2 points in case of a correct go trial.

Inter-trial-intervals (ITI, gray background) were randomly drawn from {1, 2, 4, 8, 16 s} with the respective proportion of trials {52%, 26%, 13%, 6%, 3%}.

### Experimental design and structure

Our chosen distribution of preparation times, inter-trial intervals, and no-go trials were determined by minimizing the variance inflation factor (VIF). VIF is the ratio of the mean estimation variance of all regression weights (preparation- and production-related regressors for each sequence) to the mean estimation variance had these regressors been estimated in isolation. Therefore, VIF quantifies the severity of multicollinearity between model regressors by providing an index of how much the variance of an estimated regression coefficient is increased because of collinearity. Large values for VIF mean that model regressors are not independent of each other, whereas a VIF of 1 means no inflation of variance. After optimizing the design, VIF was on average 1.15, indicating that we could separate planning and execution-related activity without a large loss of experimental power.

Participants underwent one fMRI session consisting of 10 functional runs and 1 anatomical scan. In an event-related design, we randomly interleaved 3 types of repeated single-finger presses involving the thumb (1), the middle (3), and the little (5) fingers (e.g., 111111 for thumb presses, Fig. 1B) and 3 types of multi-finger sequences (e.g., 135315). The day before the fMRI scan, participants familiarized themselves with the experimental apparatus and the go/no-go paradigm in a short behavioral session of practice outside the scanner (5 blocks, about 15-30 minutes in total). For the behavioral practice, inter-trial intervals were kept to a fixed 1 s to speed up the task, and participants were presented with different sequences from what they would see while in the scanner. These 6-item sequences were randomly selected from a pool of all possible permutations of the numbers 1, 3, and 5, with the exclusion of sequences that contained consecutive repetitions of the same number. Each sequence trial type (e.g., 111111) was repeated 5 times (2 no-go and 3 go trials), totaling 30 trials per functional run. Two periods of 10 seconds rest were added at the beginning and at the end of each functional run to allow for signal relaxation and provide a better estimate of baseline activation. Each functional run took about 5.5 minutes, and the entire scanning session (including the anatomical scan and setup time) lasted for about 75 minutes.

### Imaging data acquisition

High-field functional magnetic resonance imaging (fMRI) data were acquired on a 7T Siemens Magnetom scanner with a 32-channel head coil at Western University (London, Ontario, Canada). The anatomical T1- weighted scan of each participant was acquired halfway through the scanning session (after the first 5 functional runs) using a Magnetization-Prepared Rapid Gradient Echo sequence (MPRAGE) with a voxel size of 0.75x0.75x0.75 mm isotropic (field of view = 208 x 157 x 110 mm, encoding direction coronal). To measure the blood-oxygen-level dependent (BOLD) responses in human participants, each functional scan (330 volumes) used the following sequence parameters: GRAPPA 3, multi-band acceleration factor 2, repetition time [TR] = 1.0 s, echo time [TE] = 20 ms, flip angle [FA] = 30 deg, slice number: 44, voxel size: 2x2x2 mm isotropic. To estimate and correct for magnetic field inhomogeneities, we also acquired a gradient echo field map with the following parameters: transversal orientation, a field of view: 210 x 210 x 160 mm, 64 slices, 2.5 mm thickness, TR = 475 ms, TE = 4.08 ms, FA = 35 deg.

### Preprocessing and univariate analysis

Preprocessing of the functional data was performed using SPM12 (fil.ion.ucl.ac.uk/spm) and custom MATLAB code. This included correction for geometric distortions using the gradient echo field map (Hutton et al., 2002), and motion realignment to the first scan in the first run (3 translations: x, y, z; 3 rotations: pitch, roll yaw). Due to the short TR, no slice timing corrections were applied. The functional data were co-registered to the anatomical scan, but no normalization to a standard template or smoothing was applied during preprocessing. To allow magnetization to reach equilibrium, the first four volumes of each functional run were discarded. The pre- processed images were analyzed with a general linear model (GLM). We defined separate regressors for each combination of the 6 finger-actions (single, multi) x 3 phases (preparation go, preparation no-go, production), resulting in a total of 18 regressors (12 go + 6 no-go), plus the intercept, for each run. We also conducted an analysis where the same preparation regressor was used in go and no-go, which resulted in qualitatively similar as reported here. For the main text, however, we decided to be conservative and not use the regressors for the preparation of go trials, thereby controlling for a residual bias from the execution-related activity onto the preceding planning-related activity. Each regressor consisted of boxcar functions of length 2s convolved with a two-gamma canonical hemodynamic response function with a peak onset at 5 s and a post-stimulus undershoot minimum at 10 s. Given the relatively low error rates (i.e., number of error trials over the total number of trials, timing errors: 7.58 ± 0. 62 %; press errors: 1.18 ± 0.26 %; see Task above), all trials were included to estimate the regression coefficients, regardless of whether the execution was correct or erroneous. Ultimately, the first-level analysis resulted in activation images (beta maps) for each of the 18 conditions per run, for each of the participants.

### Surface reconstruction

Based on the 0.75mm anatomical scan, we reconstructed each individual cortical surface. Individual participants’ cortical surfaces were reconstructed using Freesurfer (Dale et al., 1999). First, we extracted the white-gray matter and pial surfaces from each participant’s anatomical image. Next, we inflated each surface into a sphere and aligned it using sulcal depth and curvature information to the Freesurfer average atlas (Fischl et al., 1999). Subsequently, surfaces were resampled to a left-right symmetric template (fs_LR.164k.spec; Van Essen et al., 2012) included in the connectome workbench distribution (Marcus et al., 2011). The functional imaging data (2mm resolution) was then mapped onto this surface. In this analysis we excluded voxels that lay within the sulci and touch both banks (with more than 25% of the voxel volume in the grey matter on both sides sulcus). The individual surface maps were brought into alignment by morphing the surfaces based on the depth and curvature (van Essen et al., 2012). This approach is currently the state-of-the-art in imaging analysis to achieve the best regional specificity of group analysis despite the considerable inter-subject variability in cortical folding.

### Regions of interest (ROI)

We focused our imaging analysis on the dorsolateral aspect of the contralateral (left) hemisphere (purple area of Fig. 2A), including the motor regions of the medial wall.

**Figure 2.**
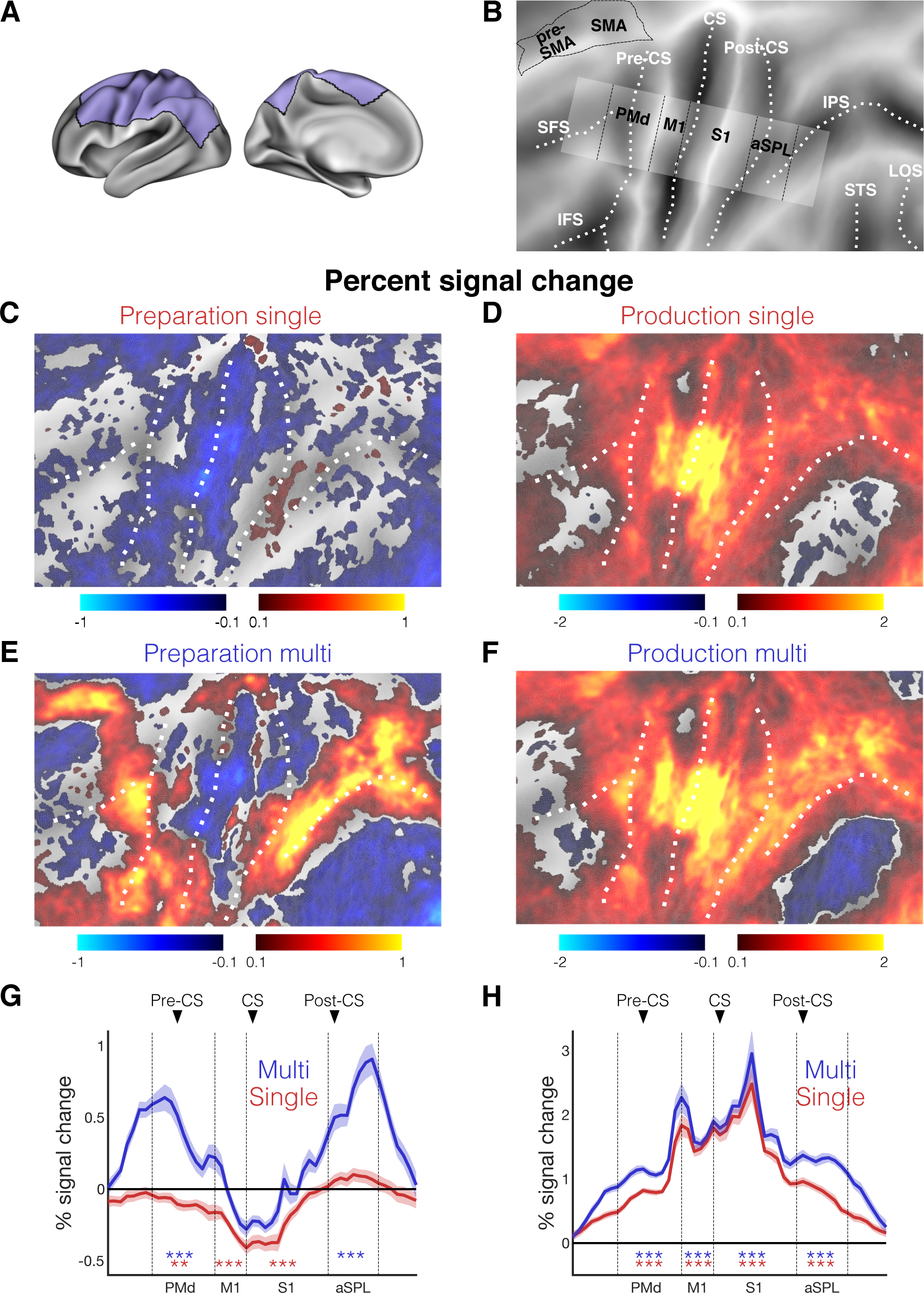
Premotor and superior parietal lobule are activated during sequence preparation. **(A)** inflated cortical surface of the contralateral (left) hemisphere, highlighting the displayed areas (B-F, purple). **(B)** Flat representation of the neocortex with major sulci indicated by white dotted lines, and the boundaries of different regions indicated by black dashed lines. The strip highlighted in white was used for the profiles (G, H) and region-of-interest definition. **(C)** Group-averaged percent signal change during preparation and **(D)** production of single-finger sequences. **(E,F)** Same as (C,D) but for multi-finger sequences. **(G)** Profile ROI analysis (see Materials and methods) of the mean percent signal change (± standard error of the mean [SEM]) during the preparation and **(H)** production of single-finger (red) and multi-finger sequences (blue). The x-axis corresponds to Brodmann areas (BA) shown in (B). **p < 0.01, ***p<0.001 in a two-sided one-sample t-test vs. zero for selected ROIs. Vertical black lines mark the approximate boundaries between the BAs (see Methods). Black triangles point to the approximate location of the main anatomical landmark. Sulci: superior frontal sulcus (SFS), inferior frontal sulcus (IFS), precentral sulcus (Pre-CS), central sulcus (CS), post central sulcus (Post-CS), intra-parietal sulcus (IPS), lateral occipital sulcus (LOS), superior temporal sulcus (STS). ROIs: anterior superior-parietal lobule (aSPL, BA 5), primary somatosensory cortex (S1, BA 3, 1, 2), primary motor cortex (M1), dorsal premotor cortex (PMd, BA 6), secondary motor area (pre-SMA and SMA, BA 6).

To summarize the results and for statistical analysis, we have used a set of motor-related ROIs on the cortical surface that we have used consistently in previous papers (Ariani et al., 2022; Berlot et al., 2020, 2019; Jörn Diedrichsen et al., 2013; Yokoi et al., 2018). The definition relies on a post-mortem cytoarchitectonic analysis of 10 human brains (Fischl et al., 2008) that were normalized into a spherical (surface-based) group atlas. In comparison to a volume-based normalization of the same maps (Eickhoff et al., 2007), this approach leads to a cleaner separation of cortical areas. One of the limitations of such ROI-based approaches is potentially combining functional heterogeneous regions into a single ROI. For the current paper, we therefore chose a more refined approach that allows us to study differences in function within these ROIs in a more continuous manner, while still summarizing the data better than the map-wise approach.

For visualization of the functional profiles along the cortical surface, we defined an anterior-posterior line through the anatomically defined hand-knob (Yousry et al., 1997) that ran approximately orthogonal to the boundaries between different Brodmann areas (BA; Fischl et al., 2008). By extending the line ±20 mm above and below the line, we defined a strip of surface area in each hemisphere that mainly captures the hand area of M1 and S1 (white area in Fig. 2B). We subdivided this area into 50 vertical sections running from anterior to posterior, allowing us to summarize the result on a one-dimensional profile-view (e.g., Fig 2G,H; Fig. 3E,F). For statistical analysis only, we combined results by major region (grouping vertical sections together) according to the cytoarchitectonic probabilistic atlas. We defined the region of interest for the primary motor cortex (M1; BA4), primary somatosensory cortex (S1; BA 1, 2, and 3), dorsal premotor cortex (PMd; BA 6), and the anterior parietal lobules (aSPL; BA 5) by selecting the strips that had the highest probability (averaged over all vertices within the strip) as determined by a probabilistic cytoarchitectonic atlas (Fischl et al., 2008) of belonging to their respective BA. The supplementary motor areas (SMA) lay outside of the defined strip of surface area. We therefore defined the SMA ROI by choosing the part of BA6 that was situated in the medial wall.

**Figure 3.**
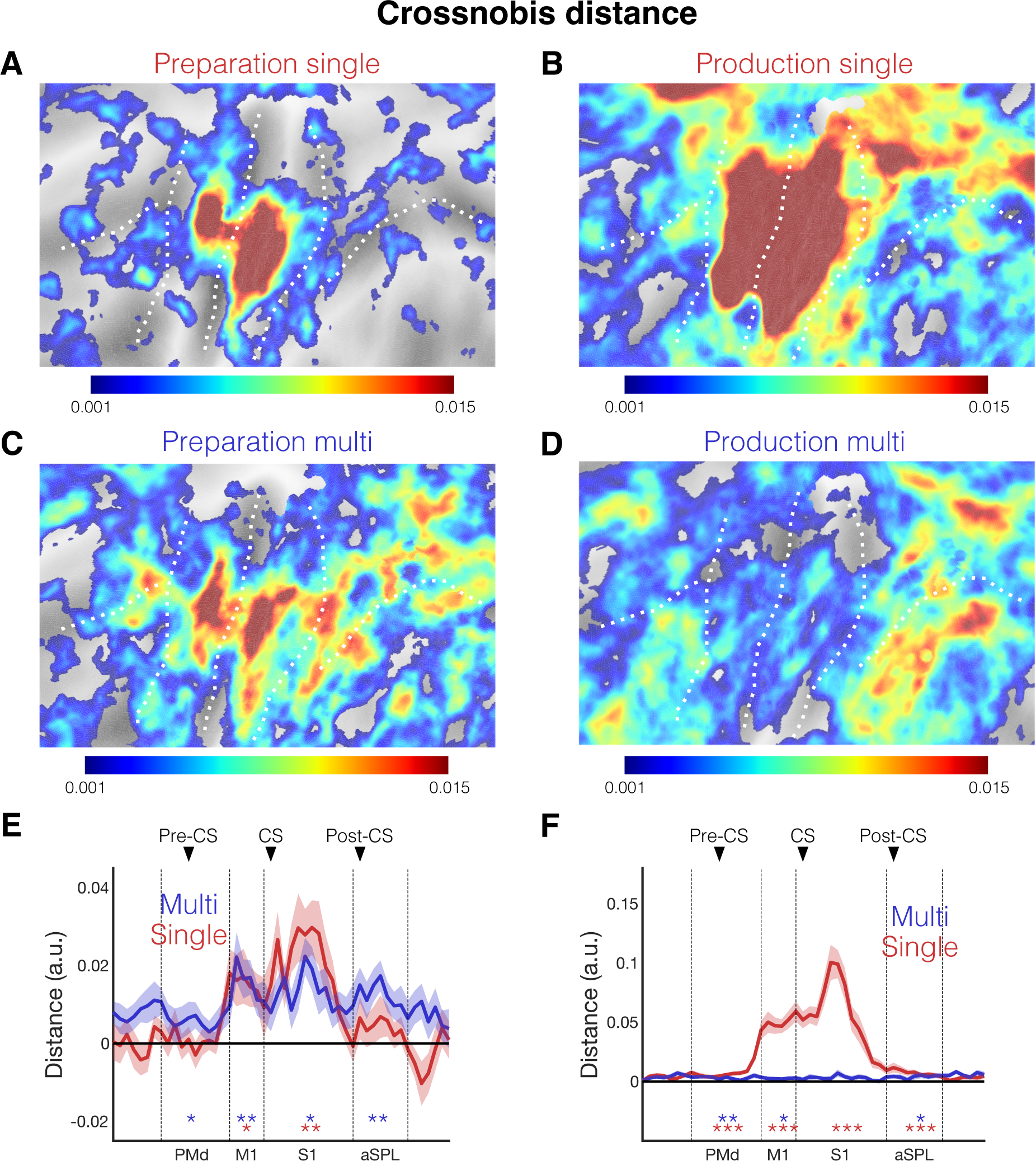
Sequence representations during preparation and production phases. **(A)** Group-averaged multivariate searchlight map of the crossnobis distance between the preparation and **(B)** production of the three single-finger. **(C)** Same as A but for mean crossnobis distance between the preparation and (**D**) production of three multi-finger sequences. **(E)** Profile ROI analysis of the mean crossnobis distance (±SEM) during the preparation of single-finger (light red line) and multi-finger (light blue) sequences. **(F)** Same as E but for the production of single-finger (dark blue) and multi-finger (dark red) sequences. *p < 0.05, **p<0.01, ***p<0.001 in a two-sided one-sample t-test vs. zero for selected ROIs.

ROI-based analyses were conducted in the space of the individual data acquisition for each individual participant by determining the voxel that would be projected onto the set of surface nodes associated with each ROI. In this analysis (as well as for surface-based mapping), we excluded voxels with more than 25% of their volume in the grey matter on the opposite side of a sulcus. This avoided cross-contamination of activity measured in M1 and S1, as well as across the pre-central and post-central sulcus. No smoothing of functional activity in the volume was applied. This approach has allowed us in one previous paper to carefully analyze the specialization of subregions of human S1 and M1 (Arbuckle et al., 2022).

### Analysis of activation

We calculated the percent signal change for each condition relative to the baseline activation for each voxel for each functional run and averaged it across runs. For ROI analysis, these values were averaged across all voxels in the native volume space of each participant selected for the respective ROI. For surface-based group maps, individual data were projected onto the group map via the individual surfaces, using all voxels touching the line that connected corresponding nodes on the pial and white-matter surfaces.

Statistical analyses to assess the cortical activity of each sequence type during each phase of preparation or production included a two-sided one-sample t-test vs. zero. For statistical tests on the surface, we used an uncorrected threshold of *p=0.001* and controlled the family-wise error by calculating the size of the largest suprathreshold cluster across the entire cortical surface (estimated smoothness of FWHM 7.9 mm) that would be expected by chance (p=0.05) using Gaussian field theory as implemented in the fmristat package (Worsley et al., 1996).

### Multivariate distance analysis

To evaluate which brain areas displayed sequence-specific representations, we used the representational similarity analysis (Kriegeskorte & Diedrichsen, 2019). We calculated the cross-validated Mahalanobis distances (Walther et al., 2016) between evoked regional patterns (beta estimates from first-level GLM) of different pairs of conditions, 6 sequences (3 single, 3 multi) x 2 phases (preparation no-go, production). Prior to calculating the distances, beta weights for each condition were spatially pre-whitened (i.e., weighted by the matrix square root of the noise covariance matrix, as estimated from the residuals of the GLM). The noise covariance matrix was slightly regularized towards a diagonal matrix (Ledoit & Wolf, 2003). Multivariate pre-whitening has been shown to increase the reliability of dissimilarity estimates (Walther et al., 2016).

Cross-validation ensures the distance estimates are unbiased, such that if two patterns differ only by measurement noise, the mean of the estimated value would be zero (Diedrichsen et al., 2020). This also means that estimates can sometimes become negative. Therefore, dissimilarities significantly larger than zero indicate that the two patterns are reliably distinct, akin to an above-chance performance in a cross-validated pattern classification analysis.

Multivariate analysis was conducted for ROI and surface analysis. For surface-based maps, we also conducted a searchlight analysis (e.g., Fig 3A). For each surface node, we selected a circular region of 100 voxels (with a maximal radius of 12 mm) and assigned the result to the central node of the searchlight.

### Dispersion metric

To determine whether there were significant differences in the dispersion of the representations for multi- and single-finger sequences during preparation, we first set all negative dissimilarity values to 0. We then normalized the dissimilarities for each sequence type such that all vertices’ dissimilarities in the white strip (Fig. 2A) summed up to 1. This defined the weight of each surface vertex. The center-of-gravity (COG) per participant was then the weighted average of the x and y coordinates of these vertices on the flat map. Then, we calculated the dispersion (*d*) around this COG by calculating the squared distance between each vertex and COG and averaging them weighted by *w_i_*.

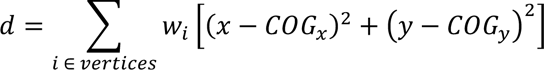

The difference in dispersion between multi- and single-finger sequences was assessed using a two-sided paired t-test.

### Correlation analysis

To assess the relationship between planning and execution-related activity, we estimated the correlation between them. For estimation, we employed two different approaches. First, we used simple Pearson’s correlation. When estimating the correlation between sequence-specific activity patterns, we first removed the mean pattern for the preparation and production phase. We then stacked the three activity patterns into a single vector. The average (across runs) activity patterns for preparation and production were correlated directly. When estimating the correlation between multi vs. single contrasts across preparation and production, we first calculated the difference between activity patterns of 3 multi-finger sequences and the corresponding single-finger movements (i.e., sequence 135351 with 111111, etc.) and then averaged the three maps. The average (across runs) difference maps for preparation and production were then correlated directly. Correlations were calculated separately within each participant. To test for a positive correlation, we performed a two-sided paired t-test against zero.

To obtain an estimate of the correlation, corrected for the level of measurement noise, we used pattern component modelling (PCM, Diedrichsen et al., 2018). The problem with Pearson’s correlations estimated on the noisy data is that they are always smaller than 1, even if the underlying patterns are identical. PCM solves this issue by modelling the noise and true pattern separately and estimating the likelihood of the data given any value of correlation (Diedrichsen et al., 2023). We created 100 correlation models with correlations in the range [0–1] in equal step sizes and assessed the likelihood of the observed data from each participant under each correlation model (Fig. 4C). First, we used the winning model as an estimate of the noise-corrected correlation (maximum-likelihood estimate). To test whether two sets of activity patterns are identical, we compared the likelihood of the best-fitting model within each participant to the likelihood of a competing correlation model (r=1), using a two-sided paired Wilcoxon signed-rank test. The choice of the best-fitting model was acquired using a cross-validated approach, estimating the group-winning model from n-1 participants, and determining the log-likelihood of this model for the left-out participant (for whom this model may not be the best one).

**Figure 4.**
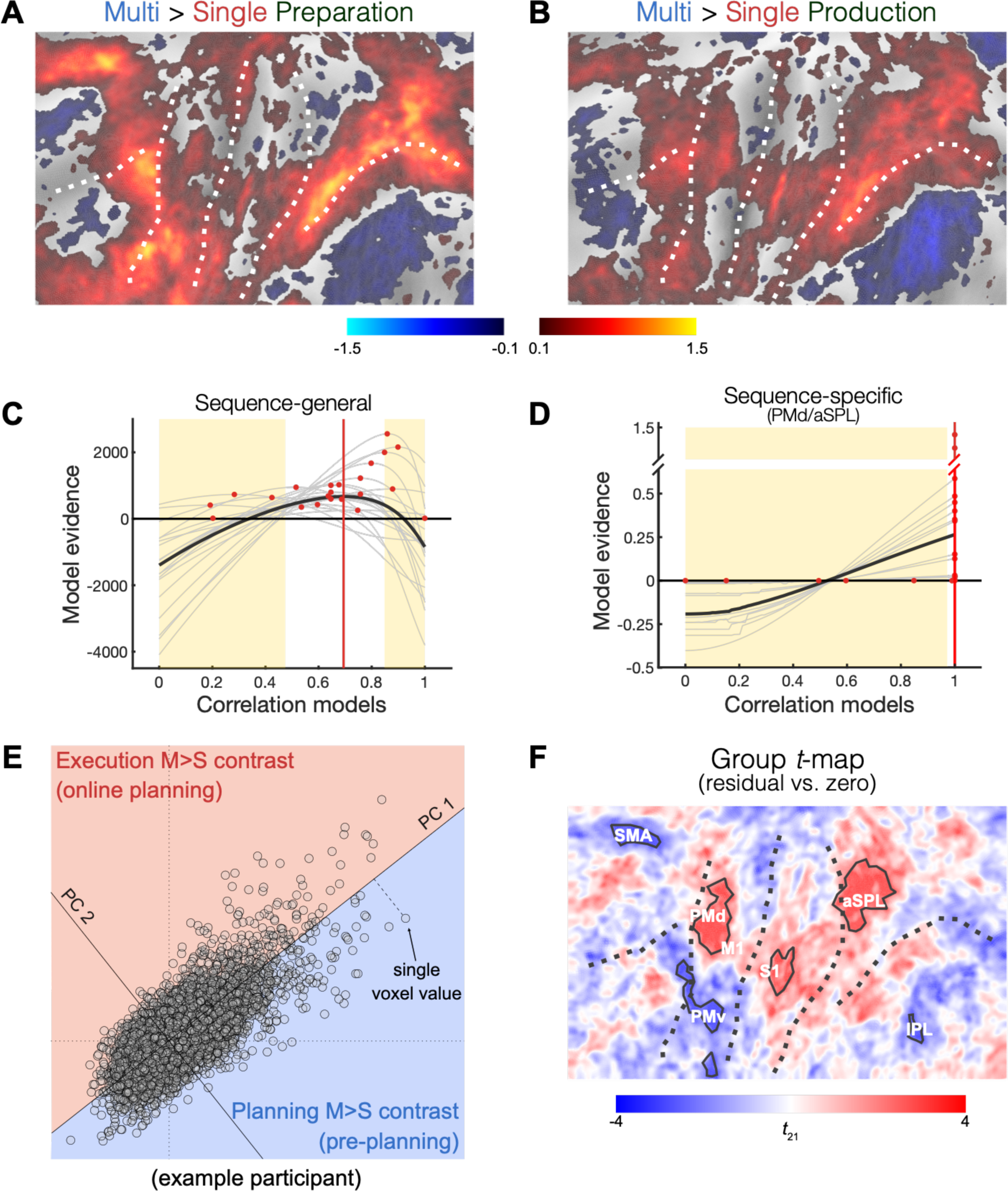
Activation maps for pre- and online planning are highly, but not perfectly, correlated. **(A)** The difference in evoked activation between multi-finger and single-finger sequences during the preparation and **(B)** production. (**C**) PCM evaluation of models assuming a correlation between the sequence-general activity patterns shown in A and B of 0 to 1. The group average (black line) and individual curves (thin gray lines) express the difference from the log-likelihood from the mean value (i.e., zero on the y-axis). Red dots indicate the best-fitting correlation model for each participant and the red solid line cross-validated best-fitting model across all participants. Yellow-shaded area indicates models that perform significantly worse than the best-fitting correlation model (p < 0.05). **(D)** Similar to (C), but for the correlation between sequence-specific activity patterns evoked during preparation and production of multi-finger sequences in PMd and aSPL. For each participant, we averaged the curve corresponding to each of the two areas. **(E)** Voxel-wise values of the preparation multi-single contrast, plotted against the voxel-wise values of the production multi-single contrast. Two principal components (PCs) of the data are shown. Voxels in the red area (positive PC2) show a preference for online planning, whereas voxels in the blue area (negative PC2) show for pre-planning. An example of a single participant is shown. **(F)** Group t-value for the second PC projected on the flat map. Areas with black solid outlines represent significant clusters. Note that for this surface-based (rather than ROI-based) analysis, there is no strict correspondence between the clusters on the map and our pre-defined ROIs. The labels next to the significant clusters correspond to the more closely matching ROI.

### Analysis of principal components

To test if the activity patterns for the difference between multi- and single-finger sequences across preparation and production are systematically different in some brain regions, we employed principle competent analysis (PCA). The simple contrast between the two difference maps could not answer this question because they have different scales. We obtained the multi vs. single contrast during preparation and production averaged across sequences for all voxels within the region of interest (Fig. 2A, purple area). For each individual, we then conducted a singular value decomposition on the 2xP matrix, in which the two rows represented the preparation and production contrasts. The first PC captured the common neural activity shared by processes happening during preparation and production while accounting for the differing degrees of activation in specific voxels. The second PC represented the tendency of each voxel to be relatively more engaged during one process than the other. We ensured that for each participant the positive value on the second PC represented relatively more engagement during the production phase. Lastly, we projected the value for each voxel to the nearest vertex and created a preference map. These maps were then submitted to a surface-based group analysis (see above).

## Results

### Pre-planning of multi-finger sequences activates the first finger in M1 and S1

First, we asked what processes occur during sequence preparation in the core sensorimotor areas S1 and M1. For single-finger movements (Fig. 2C), BOLD activity in these regions was suppressed relative to rest (Fig. 2G; M1: *t_21_=−6.939, p=7.4e−07*; S1: *t_21_=−5.508, p=1.8e−05*). For multi-finger movements (Fig. 2E), we also observed some suppression deep in the central sulcus, however, averaged across the ROI, the activation was not different from the resting baseline (M1: *t_21_=−0.692, p=0.496*; S1: *t_21_=−0.523, p=0.606*).

We then used multi-voxel pattern analysis to examine whether the fine-grained pattern of activity in these areas differed between the planned sequences. To this end, we calculated cross-validated Mahalanobis distances (Diedrichsen et al., 2020; Walther et al., 2016) between the activity patterns associated with the 3 sequences. Systematically positive values of this measure indicate reliable multivariate pattern differences. As previously reported (Ariani et al., 2022), we found that during the preparation of single-finger movements (Fig. 3A), the core hand areas of M1 and S1 exhibited finger-specific activity patterns (M1: *t_21_=2.343, p=0.029*; S1: *t_21_=3.137, p=0.005*). Analysis of multi-finger sequences revealed a similar result: although M1 and S1 were not activated overall, there were significant differences between activity patterns for three sequences (Fig. 3E; M1: *t_21_=2.991, p=0.007*; S1: *t_21_=2.829, p=0.010*).

What do these activity differences for multi-finger sequences reflect? One possibility is that during sequence preparation, the first element of the sequences is pre-activated in M1. Because each sequence started with a different finger, this would cause large differences between the activity patterns without constituting a sequence representation. This idea predicts that the activity pattern during preparation should correlate with the activity pattern observed when only the first finger movement is pressed. To test this, we correlated the preparation activity of 3 multi-finger sequences with the production activity of the corresponding single-finger sequences (i.e., sequence 135351 with 111111, etc.) after subtracting the mean activity pattern. This analysis revealed a significant correlation in both M1 (average *r=0.043, t_21_=2.366, p=0.027*) and S1 (*r=0.047, t_21_=2.285, p=0.032*) but not in PMd (*t_21_=-1.227, p=0.233*) and aSPL (*t_21_=-0.242, p=0.811*) with the values of correlation being significantly larger in M1/S1 than PMd/aSPL (*t_21_=3.068, p=0.005)*. While these correlations are very small, correlation estimated on noisy data consistently underestimate the true correlation (Diedrichsen et al., 2023). To account for measurement noise in a principled way, we used Pattern Component Modeling (PCM, see Methods) to build potential models of the correlation between activity patterns for the preparation of multi-finger sequences and production of single-finger sequences and evaluate the likelihood of the data given each model. We removed the average activity pattern common to all sequences from the patterns (separately for multi- and single-finger sequences), and then correlated the patterns across the two conditions. The group maximum-likelihood estimates of correlation indicated substantial correspondence with r=0.40 for S1 and r=0.34 for M1. Thus, there is substantial overlap between the patterns during pre-planning of multi-finger sequences and the patterns during the execution of the first finger in the sequence.

In previous studies, we have found that the activity patterns in M1 and S1 (averaged over preparation and production phases) can be explained by a temporal summation of the patterns related to the individual finger presses, with an especially high weight from the first finger in the sequence (Berlot et al., 2021; Yokoi et al., 2018). If this weighting is due to the pre-activation of the first finger during movement preparation, we would predict that the pattern differences between sequences should disappear during the execution phase, as all three involved the same three fingers (albeit in a different order). Indeed, the pattern differences were significantly attenuated during the production phase (Fig. 3D, planning vs. execution in M1: *t_21_=2.123, p=0.0458*, and S1: *t_21_=2.305, p=0.0345*), with a small pattern difference remaining only in M1 (*t_21_=2.814, p=0.0104*).

In sum, these results are consistent with the idea that the first finger in a sequence is pre-activated during the preparation phase and that after production starts, the activity pattern in M1 and S1 are determined by a temporal summation of the patterns related to the individual finger presses (Yokoi et al., 2018).

### Pre- and online planning engage a highly overlapping set of cortical areas

If the activity patterns in the M1 and S1 are related to the individual finger presses, then continuous input from higher-level regions that retain a representation of the entire sequence is required to move reliably from finger to finger. We refer to this process as planning, whether it occurs before (pre-planning) or after movement onset (online planning; Ariani & Diedrichsen, 2019; Ariani et al., 2021). Here investigate whether pre- and online planning engage exactly the same, or different sets of cortical areas. Given that the number, speed, and force of the finger presses were closely matched across single and multi-finger movements (Table 1), we assumed that both conditions involve similar execution-related processes. Therefore, the difference between the production of single-finger and multi-finger sequences should mostly reflect activity related to online planning. If pre-planning and online planning involve the exact same cortical areas, then this difference should be similar to the difference between single-finger and multi-finger movement during preparation (reflecting the need for increased pre-planning).

**Table 1.** Average execution time (ET, i.e., time needed to complete a sequence) and average peak force for single-finger and multi-finger sequences across participants. A two-sided paired t-test for a difference between conditions is reported in the last column.

|  | Single-finger | Multi-finger | Difference |
| --- | --- | --- | --- |
| ET [ms] | $1450 \pm 58$ | $1370 \pm 70$ | $t_{21}=1.815$, $p=0.083$ |
| Force [N] | $1.7 \pm 0.1$ | $1.8 \pm 0.1$ | $t_{21}=1.423$, $p=0.169$ |

In contrast to single-finger movements (Fig. 2C), preparation of multi-finger sequences (Fig. 2E) strongly activated the dorsal premotor cortex (Fig. 2G; PMd, *t_21_=5.125, p=4.4e-05*), the supplementary motor areas (SMA/pre-SMA, *t_21_=4.016, p=2.3e-04*) and the anterior part of the superior parietal lobule (aSPL, *t_21_=7.482, p=2.3e-07*). Thus, the demands of planning multiple different finger movements evoked significant brain activation in premotor and parietal areas.

During the production, we found widespread activity in both primary sensorimotor, parietal, and pre-motor areas for both sequence types (Fig. 2D, F). The contrast between these two conditions revealed higher activity for multi-finger over single-finger sequences in PMd (Fig. 2H, 4B; *t_21_=6.022, p=5.6e-6*) and aSPL (*t_21_=8.264, p=4.8e-08*). In contrast, in caudal M1 and rostral S1 the same two conditions elicited very similar activity levels, consistent with the idea that the basic motor requirements were well matched between single- and multi-finger sequences.

Importantly, the spatial distribution of multi vs. single contrast during production was very similar to multi vs. single contrast during preparation (Fig. 4A, B). When we correlated the unsmoothed voxel-wise activity maps within each participant, we found a highly significant correlation between the brain activity patterns under these two conditions (r=0.12, *t_21_=10.796, p=4.9e-10*). Thus, pre- and online planning activated overlapping cortical areas.

### Preferential activation for planning before and during movement

The strong overlap between the activity maps for pre- and online planning, however, does not tell us whether the two maps were identical (in other words, pre- and online planning activated - within each participant - exactly the same voxels) or whether there were true differences between the maps. This is because measurement noise will lead to an observed correlation smaller than 1, even if two maps are identical (Diedrichsen et al., 2023). Again, we applied PCM to model the correlation between multi-single contrast activity patterns in both preparation and production, followed by an assessment of data likelihood for each model (see Fig. 4C). The group maximum-likelihood estimate of correlation was r=0.65 with strong evidence that the patterns were not identical (against 1- corr model: *z = 3.815, p=0.0001*).

Therefore, we asked whether there were systematic differences across participants, which would indicate that some areas have a preference toward either pre- or online planning. Because the contrast during pre-planning was generally larger than during execution, we could not simply subtract the two difference maps. Instead, we plotted the multi-single difference for each voxel during preparation (Fig. 4E, x-axis) against the difference during production (y-axis). We then estimated the average relationship between the two contrasts using Principal Component Analysis (PCA) within each participant. The first principal component (PC1) captured the tendency of voxels to be similarly responsive for pre- or online planning. The loading on the second principal component captured the deviation from this lawful relationship, with positive values indicating relatively more activity during online planning and negative values more activity during pre-planning. By mapping the voxel preferences back to the surface, we created an average preference map across participants within our window of interest (Fig. 4F). This map revealed that clusters (solid outlines in Fig. 4F) in PMd (*p=5.9e-07*, corrected for multiple comparisons, see Methods), M1/S1 (*p=8.8e-06*), and aSPL (*p=5.9e-07*) were more active during online planning, while clusters in SMA (*p=3.1e-06*), the ventral premotor cortex (PMv, *p=7.2e-07* and *p=0.0096*), and the inferior parietal lobule (IPL, *p=0.0492*) were relatively more active during pre-planning (Fig. 4F). This set of results indicates that pre-planning and online planning of movement sequences preferentially activated slightly different sets of cortical areas.

### PMd and SPL maintain sequence-specific representations both during preparation and production

If the increased activation for multi-finger sequences in PMd and SPL (Fig. 4A-B) was related to pre-planning and online planning, then these areas should exhibit sequence-specific representations during both the preparation and production phases.

Indeed, for the preparation phase multivariate analysis (Fig. 3C) revealed that both areas showed sequence-specific representations of multi-finger sequences. We found significant pattern differences in PMd (Fig. 3E, *t_21_ = 2.266, p=0.034*) and aSPL (*t_21_=3.491, p=0.002*). In contrast to the preparation of single-finger movements (Fig. 3A), the representations of multi-finger sequences appeared to be more widespread on the cortical sheet. To quantify this observation, we calculated a spatial dispersion metric, which reflects the spatial variance of pattern dissimilarities (see Methods). This analysis confirmed that the planning of multi-finger sequences was associated with a more widespread representation across the sensory-motor network compared with single-finger sequences (*t_21_=3.542, p=0.002*). Therefore, premotor and parietal areas were not only more active during the preparation of multi-than single-finger sequences (Fig. 4A), but also represented the identity of the sequence.

Importantly, we also found that these representations were maintained during the execution phase (Fig. 3D, PMd: *t_21_=3.221, p=0.004*; aSPL: *t_21_=2.490, p=0.021*). When comparing the multivariate distances between preparation (Fig. 3C) and execution (Fig. 3D) we found a strong reduction in M1 (*t_21_=2.123, p=0.045*) and S1 (*t_21_=2.305, p=0.0314*). In contrast, the pattern differences in PMd and aSPL did not attenuate from the preparation to the execution phase (PMd: *t_21_=0.271, p=0.7889*, and aSPL: *t_21_=1.895, p=0.0719*). Note that the multivariate distances reported in Fig. 3 are measured with noise, so the small-scale differences that can be observed between Fig. 3C and Fig. 3D should not be overinterpreted. When conducting a map-wise comparison, none of the subtle differences are significant after controlling for multiple comparisons. We therefore chose a ROI approach to increase our statistical power. These results are consistent with the idea that premotor and parietal areas are involved in the planning of movement sequences both before and during sequence production.

Finally, we asked whether the sequence-specific representations in these areas remained stable or whether they changed dynamically from preparation to production. To address this question, we again used PCM. We removed the average activity pattern common to all multi-finger sequences from the patterns (separately for preparation and production phase), and then correlated these sequence-specific pattern differences between sequences across the two phases. Figure 4D shows the likelihood curves for different correlation models averaged across PMd and aSPL. The small signal-to-noise for relatively subtle differences between multi-finger sequences resulted in small model evidence for almost all participants. The cross-validated group-winning model predicted that sequence-specific activity patterns were perfectly correlated with strong evidence that patterns were not uncorrelated (against 0-corr model: PMd: *z = 2.646, p=0.008*; aSPL: *z = 2.873, p=0.004*). These results suggest that sequence-specific representations remain stable across pre-planning and execution.

## Discussion

We used 7T fMRI and multivariate analyses to investigate the role of human cortical areas in the preparation of movement sequences, separating processes that occur before and during movement production. We found that primary sensorimotor cortices (M1, S1) showed activity patterns resembling the first movement in the sequence during preparation, and of a temporal summation of individual movements during production. In contrast, secondary sensorimotor areas (PMd, SMA, SPL) were more activated during both pre- and online planning of motor sequences. These regions also maintained a stable representation of the sequence across preparation and production phases.

In previous fMRI studies, we found that the activity patterns in M1 and S1 during sequence production can be explained by the linear combination of the patterns associated with the individual movements (Berlot et al., 2021; Yokoi & Diedrichsen, 2019). The pattern for the first finger in the sequence contributed substantially more to the overall pattern (average over preparation and execution) than all the subsequent fingers (Yokoi et al., 2018). Our results now provide evidence that this *‘first finger effect’* was caused by the pre-activation of the first sequence element during preparation rather than by the transition from planning to execution state space (Kaufman et al., 2016; Yokoi et al., 2018): the pre-planning of a multi-finger sequence activated a pattern similar to that observed during the execution of the first movement in that sequence.

In a study using sequences of object-directed reach-to-grasp movements, Gallivan et al. (2016) found that the activity patterns in M1 were also slightly different between two sequences that started with the same movement but differed in the second movement. This suggests that, not only the first, but possibly also the second movement may be reflected in M1 preparatory activity patterns. This observation would be consistent with the “competitive queuing” hypothesis (Averbeck et al., 2002), the idea that the first movement in a sequence being most, the second less, and subsequent movements even less activated (Kornysheva et al., 2019). However, because the first reach-to-grasp movement was always the same across conditions, it was unclear whether these findings would generalize to longer and more complex sequences where every sequence element is different and needs to be planned anew from trial to trial.

After sequence production began, the pattern differences in M1 and S1 between the 3 multi-finger sequences were strongly attenuated. This is likely due to the fact the pattern corresponding to the single-finger movements were sequentially activated. Due to the low temporal resolution of fMRI, these patterns combined additively, such that the three multi-finger sequences (which were matched in terms of the involved fingers) elicited overall similar activity pattern when we isolated the activity during the production phase.

These results also suggest that the production of the sequence cannot be maintained autonomously by primary motor areas, but that it depends on input from secondary motor areas (Russo et al., 2020; Tanji & Shima, 1994; Yokoi & Diedrichsen, 2019). Consistent with this idea, we found that PMd, SMA, and SPL (but not caudal M1 and rostral S1) were more activated during the preparation and production of multi-finger as compared to single-finger movements. We also found sequence-specific activity patterns in these areas, which were correlated across preparation and production phases. This suggests that premotor and parietal areas maintained a stable representation of the sequence throughout, likely to drive the next movements in M1.

As shown in recent behavioral studies (Ariani & Diedrichsen, 2019; Ariani et al., 2021; Kashefi et al., 2023), even relatively short 5-item sequences cannot be fully pre-planned. Instead, planning of remaining items needs to continue throughout sequence production. To determine if the same areas are involved in the pre- and online planning, we contrasted the univariate multi-finger to single-finger activity maps. As basic execution processes were matched, we interpreted these contrasts as reflecting the increased need for planning. We found that the contrast maps were highly correlated across preparation and production phases (Fig. 4C), indicating that pre-planning and online planning activated similar regions. Within two of these regions, the PMd and aSPL, we also found that the sequence-specific activity patterns were highly correlated across these preparation and production (Fig. 4D), suggesting that the sequence-specific representations remained stable. Together these two findings are evidence that the processes involved in online planning are similar to those involved in pre-planning. Despite considerable overlap in brain activation between pre- and online planning (Fig. 3A,B), however, further analysis revealed small but systematic differences in the involvement of different brain areas: PMd, S1, and SPL tended to be more active during online planning, and SMA and ventral premotor cortex (PMv) more during pre-planning. This indicates that the two processes are not exactly identical. One possible explanation could be the differential engagement of brain areas in memory encoding and memory retrieval processes that, as part of planning processes, happened during preparation and production, respectively. The exact nature of these differences, however, remains to be tested in future studies.

Our results offer testable predictions for future neurophysiological recordings in sensorimotor areas of non-human primates. They predict that M1 shows fast dynamics related to each individual movement, with linear superposition of subsequent movements (Zimnik & Churchland, 2021). Additionally, they predict the presence of sequence-specific representation in premotor and superior parietal areas both during preparation and production. The observed consistency of these representations during preparation and production suggests that these representations have slow dynamics and change very little from preparation to production. However, it is also possible that these patterns are stable at a voxel level—i.e., pre- and online planning activate the same cortical columns—but show faster dynamics at the single neuron level. Studying these representations at a neuronal level at high temporal resolution will provide additional insight into how the motor system solves the fundamental problem of serial order in behavior (Lashley, 1951).

## Author contributions

G.A. and J.D. designed research; G.A. performed research; M.S. and G.A. analyzed data; All authors drafted and edited the manuscript.

## Acknowledgments

This work was supported by a Discovery Grant from the Natural Sciences and Engineering Research Council of Canada (NSERC, RGPIN-2016-04890) and a project grant from the Canadian Institutes of Health Research (CIHR, PJT-175010) to J.D., and the Canada First Research Excellence Fund (BrainsCAN).

## References

Arbuckle, S. A., Andrew Pruszynski, J., & Diedrichsen, J. (2022). Mapping the Integration of Sensory Information across Fingers in Human Sensorimotor Cortex. The Journal of Neuroscience: The Official Journal of the Society for Neuroscience, 42(26), 5173–5185.

Ariani, G., & Diedrichsen, J. (2019). Sequence learning is driven by improvements in motor planning. Journal of Neurophysiology, 121(6), 2088–2100.

Ariani, G., Kordjazi, N., Pruszynski, J. A., & Diedrichsen, J. (2021). The Planning Horizon for Movement Sequences. ENeuro, 8(2). 10.1523/ENEURO.0085-21.2021

Ariani, G., Oosterhof, N. N., & Lingnau, A. (2018). Time-resolved decoding of planned delayed and immediate prehension movements. Cortex; a Journal Devoted to the Study of the Nervous System and Behavior, 99, 330–345.

Ariani, G., Pruszynski, J. A., & Diedrichsen, J. (2022). Motor planning brings human primary somatosensory cortex into action-specific preparatory states. ELife, 11. 10.7554/eLife.69517

Averbeck, B. B., Chafee, M. V., Crowe, D. A., & Georgopoulos, A. P. (2002). Parallel processing of serial movements in prefrontal cortex. Proceedings of the National Academy of Sciences of the United States of America, 99(20), 13172–13177.

Berlot, E., Popp, N. J., & Diedrichsen, J. (2020). A critical re-evaluation of fMRI signatures of motor sequence learning. ELife, 9, e55241.

Berlot, E., Popp, N. J., Grafton, S. T., & Diedrichsen, J. (2021). Combining Repetition Suppression and Pattern Analysis Provides New Insights into the Role of M1 and Parietal Areas in Skilled Sequential Actions. The Journal of Neuroscience: The Official Journal of the Society for Neuroscience, 41(36), 7649–7661.

Berlot, E., Prichard, G., O’Reilly, J., Ejaz, N., & Diedrichsen, J. (2019). Ipsilateral finger representations in the sensorimotor cortex are driven by active movement processes, not passive sensory input. Journal of Neurophysiology, 121(2), 418–426.

Dale, A. M., Fischl, B., & Sereno, M. I. (1999). Cortical surface-based analysis. I. Segmentation and surface reconstruction. NeuroImage, 9(2), 179–194.

Diedrichsen, J., Shahbazi, M., Ariani, G., & Berlot, E. (2023). Estimating correlations between noisy activity patterns. A tricky problem with a generative solution. *Diedrichsenlab*. https://www.diedrichsenlab.org/BrainDataScience/noisy_correlation/index.htm

Diedrichsen, Jörn, Berlot, E., Mur, M., Schütt, H. H., Shahbazi, M., & Kriegeskorte, N. (2020). Comparing representational geometries using whitened unbiased-distance-matrix similarity. In *arXiv [stat.AP]*. arXiv. http://arxiv.org/abs/2007.02789

Diedrichsen, Jörn, Wiestler, T., & Krakauer, J. W. (2013). Two distinct ipsilateral cortical representations for individuated finger movements. Cerebral Cortex, 23(6), 1362–1377.

Diedrichsen, Jörn, Yokoi, A., & Arbuckle, S. A. (2018). Pattern component modeling: A flexible approach for understanding the representational structure of brain activity patterns. NeuroImage, 180, 119–133.

Eickhoff, S. B., Grefkes, C., Zilles, K., & Fink, G. R. (2007). The somatotopic organization of cytoarchitectonic areas on the human parietal operculum. *Cerebral Cortex (New York*, N.Y.: 1991), *17*(8), 1800–1811.

Fischl, B., Sereno, M. I., Tootell, R. B., & Dale, A. M. (1999). High-resolution intersubject averaging and a coordinate system for the cortical surface. Human Brain Mapping, 8(4), 272–284.

Fischl, Bruce, Rajendran, N., Busa, E., Augustinack, J., Hinds, O., Yeo, B. T. T., Mohlberg, H., Amunts, K., & Zilles, K. (2008). Cortical folding patterns and predicting cytoarchitecture. Cerebral Cortex, 18(8), 1973– 1980.

Gallivan, J. P., Johnsrude, I. S., & Flanagan, J. R. (2016). Planning Ahead: Object-Directed Sequential Actions Decoded from Human Frontoparietal and Occipitotemporal Networks. Cerebral Cortex, 26(2), 708–730.

Hutton, C., Bork, A., Josephs, O., Deichmann, R., Ashburner, J., & Turner, R. (2002). Image distortion correction in fMRI: A quantitative evaluation. NeuroImage, 16(1), 217–240.

Kashefi, M., Reschechtko, S., Ariani, G., Shahbazi, M., Die-drichsen, J., & Andrew Pruszynski, J. (2023). Interaction of multiple future movement plans in sequential reaching. In bioRxiv (p. 2023.05.24.542099). 10.1101/2023.05.24.542099

Kaufman, M. T., Seely, J. S., Sussillo, D., Ryu, S. I., Shenoy, K. V., & Churchland, M. M. (2016). The largest response component in the motor cortex reflects movement timing but not movement type. ENeuro, 3(4), ENEURO.0085-16.2016.

Kornysheva, K., Bush, D., Meyer, S. S., Sadnicka, A., Barnes, G., & Burgess, N. (2019). Neural Competitive Queuing of Ordinal Structure Underlies Skilled Sequential Action. Neuron, 101(6), 1166–1180.e3.

Kriegeskorte, N., & Diedrichsen, J. (2019). Peeling the Onion of Brain Representations. Annual Review of Neuroscience, 42, 407–432.

Lashley, K. S. (1951). The Problem of Serial Order in Behavior.

Ledoit, O., & Wolf, M. N. (2003). Honey, I shrunk the sample covariance matrix. SSRN Electronic Journal. 10.2139/ssrn.433840

Marcus, D. S., Harwell, J., Olsen, T., Hodge, M., Glasser, M. F., Prior, F., Jenkinson, M., Laumann, T., Curtiss, S. W., & Van Essen, D. C. (2011). Informatics and data mining tools and strategies for the human connectome project. Frontiers in Neuroinformatics, 5, 4.

Russo, A. A., Khajeh, R., Bittner, S. R., Perkins, S. M., Cunningham, J. P., Abbott, L. F., & Churchland, M. M. (2020). Neural Trajectories in the Supplementary Motor Area and Motor Cortex Exhibit Distinct Geometries, Compatible with Different Classes of Computation. Neuron, 107(4), 745–758.e6.

Shima, K., Isoda, M., Mushiake, H., & Tanji, J. (2006). Categorization of behavioural sequences in the prefrontal cortex. Nature, 445(7125), 315–318.

Tanji, J., & Shima, K. (1994). Role for supplementary motor area cells in planning several movements ahead. Nature, 371(6496), 413–416.

Van Essen, D. C., Glasser, M. F., Dierker, D. L., Harwell, J., & Coalson, T. (2012). Parcellations and hemispheric asymmetries of human cerebral cortex analyzed on surface-based atlases. Cerebral Cortex, 22(10), 2241– 2262.

Walther, A., Nili, H., Ejaz, N., Alink, A., Kriegeskorte, N., & Diedrichsen, J. (2016). Reliability of dissimilarity measures for multi-voxel pattern analysis. NeuroImage, 137, 188–200.

Worsley, K. J., Marrett, S., Neelin, P., Vandal, A. C., Friston, K. J., & Evans, A. C. (1996). A unified statistical approach for determining significant signals in images of cerebral activation. Human Brain Mapping, 4(1), 58–73.

Yokoi, A., Arbuckle, S. A., & Diedrichsen, J. (2018). The Role of Human Primary Motor Cortex in the Production of Skilled Finger Sequences. The Journal of Neuroscience: The Official Journal of the Society for Neuroscience, 38(6), 1430–1442.

Yokoi, A., & Diedrichsen, J. (2019). Neural Organization of Hierarchical Motor Sequence Representations in the Human Neocortex. Neuron, 103(6), 1178–1190.e7.

Yousry, T. A., Schmid, U. D., Alkadhi, H., Schmidt, D., Peraud, A., Buettner, A., & Winkler, P. (1997). Localization of the motor hand area to a knob on the precentral gyrus. A new landmark. Brain: A Journal of Neurology, 120 ( *Pt 1*), 141–157.

Zimnik, A. J., & Churchland, M. M. (2021). Independent generation of sequence elements by motor cortex. Nature Neuroscience, 24(3), 412–424.

